# Impacts of Environmental Nickel and Chromium Exposure on the Blood Proteome of House Sparrows (*Passer domesticus*) in New Caledonia

**DOI:** 10.64898/2026.11.08.764441

**Authors:** Zeshan Ali, Riccardo Ton, Max Mclennan-Gillings, Mark Patrick Taylor, Simon Griffith, Paul A. Haynes

**Affiliations:** School of Natural Sciences, Macquarie University, Sydney, NSW, 2109, Australia; Cavanilles Institute of Biodiversity and Evolutionary Biology, University of Valencia, Valencia, 46980, Spain; Environment Protection Authority Victoria, Centre for Applied Sciences, Melbourne, Victoria, 3085, Australia

## Abstract

New Caledonia holds some of the world’s largest ultramafic mineral deposits, and nickel (Ni) smelting operations have created a gradient of Ni and chromium (Cr) contamination across the archipelago. Molecular impacts of Ni and Cr toxicity on wildlife remain largely unexplored. Here, we used the house sparrow (*Passer domesticus*) as a sentinel species to characterise the blood proteomic response to Ni and Cr exposure near the Doniambo smelter in Nouméa. Blood was collected from 52 free-living sparrows across 18 sites classified by soil metal concentration. Plasma and blood cell pellet fractions were analysed by liquid chromatography-tandem mass spectrometry (LC-MS/MS). Plasma proteomics identified histidine-rich glycoprotein (HRG) as the most prominent responder, with approximately 2.1-fold elevation in exposed relative to low-exposure birds. HRG contains a conserved histidine-rich region that binds divalent cations including Ni²⁺, and paired plasma-liver analysis confirmed hepatic-driven synthesis (R² = 0.36, p = 0.008). Blood cell pellet proteomics revealed that Ni-Cr co-exposure predominantly disrupts the nuclear compartment: histone depletion indicated chromatin collapse, NUP153 dysregulation impaired nuclear pore function, and VRK1/TRIP13 upregulation suggested DNA damage response activation coupled with mitotic checkpoint override. Concurrent RNA processing collapse (DDX17, DHX15, RBMX, ALYREF) and changes in telomere-associated proteins (TERF2IP, hnRNPA1 and VRK1) implicated disrupted genome maintenance. These findings identify HRG as a candidate plasma biomarker of Ni-Cr exposure and reveal a coordinated nuclear toxicity signature linking heavy metal contamination to DNA damage and epigenetic disruption in a free-living vertebrate.

## 1. Introduction

New Caledonia holds about 7% of the world’s nickel reserves and is one of the largest global nickel producers [1]. Approximately one-third of the main island is covered by highly erodible ultramafic formations which are characterised by high concentrations of heavy metals, including nickel (Ni), chromium (Cr), cobalt (Co), and manganese (Mn), as well as low levels of essential nutrients such as phosphorus (P), potassium (K), and calcium (Ca) [2, 3][4]. The combination of erosion and intensive mining activities has accelerated the transfer of these heavy metals from ultramafic soil into surrounding environmental compartments, including air, water, and vegetation. As a result, humans and wildlife in the mining and ultramafic areas are exposed to high levels of Ni and Cr, among other heavy metals [4–8], raising concerns about potential health impacts in both the wildlife and human populations.

While some heavy metals, such as zinc (Zn), copper (Cu), manganese (Mn), cobalt (Co), and chromium (Cr), are required in trace amounts in living systems, elevated concentrations can be toxic [9]. Nickel and chromium are of particular concern due to their toxicity in humans. Ni disrupts metal-dependent enzymes by displacing iron (Fe) and Zn [10, 11], induces reactive oxygen species (ROS), and damages DNA through impaired repair mechanisms [12, 13]. These effects may lead to various types of cancer, respiratory disease, and neurotoxicity in mammals [14, 15]. Similarly, Cr (VI) is a known carcinogen and neurotoxin [16, 17]. Evidence of heavy metal toxicity in New Caledonian ecosystems is emerging: a recent study of marbled eels (*Anguilla marmorata*) downstream of mining operations documented significant accumulation of Ni, Cr, Co, and Mn in liver, kidney, and gill tissues, accompanied by mitochondrial dysfunction, oxidative stress, and immunotoxicity [5]. These findings suggest that chronic metal exposure compromises physiological function in aquatic species and may be driving population declines in contaminated rivers.

New Caledonia is home to a unique biodiversity, and many endemic species are in decline due to anthropogenic pressures [18]. Understanding how heavy metal contamination affects wildlife is essential for conservation efforts, yet the physiological impacts on most taxa remain poorly characterized. Avian species offer advantages as bioindicators: their high metabolic rates, diverse feeding behaviours, and frequent contact with contaminated substrates make them highly susceptible to environmental toxicants while also reflecting exposure across multiple environmental compartments [19, 20]. Moreover, birds integrate exposure over time through accumulated body burdens, providing insights into chronic toxicity that complement short-term measurements from water or soil sampling.

Among potential avian species, the house sparrow (*Passer domesticus*) is particularly well-suited for toxicological studies in New Caledonia. Introduced in the late nineteenth century, this species is now widely distributed across habitats with elevated heavy metal concentrations, including urban areas adjacent to mining activities. House sparrows have proven effective for monitoring heavy metal pollution globally [19, 21, 22], and their broad ecological plasticity enables population persistence across diverse environmental conditions. Despite their utility as sentinel species, the molecular mechanisms underlying heavy metal toxicity in house sparrows, and avian species more broadly, remain poorly understood. Investigating these pathways is essential for linking environmental exposure to physiological outcomes and informing conservation strategies in metal-contaminated landscapes.

To elucidate the molecular mechanisms of heavy metal toxicity in avian species, we employed house sparrows as our model organism and applied mass spectrometry-based proteomics. Proteomics offers distinct advantages over traditional toxicological endpoints by enabling system-wide quantification of protein abundance changes. Blood is an ideal biofluid for toxicological studies due to its easy accessibility through minimally invasive sampling, and it provides a comprehensive snapshot of systemic physiological changes [23]. However, plasma proteomics is analytically challenging due to the dominance of highly abundant proteins, which constitute ∼ 99% of the total plasma protein mass, thereby obscuring the detection of lower-abundance biomarkers [24]. Moreover, the plasma proteome may overlook biologically relevant proteins present in the cellular compartment, including red blood and immune cells. To address these limitations, we analysed both plasma and blood cell pellets, employing an abundant protein depletion strategy for the cellular fraction.

In this study, we collected blood samples from house sparrows across sites in New Caledonia with varying soil concentrations of heavy metals, which were categorized into low, medium, and high contamination groups based on Ni and Cr levels. To assess the proteome response to heavy metal exposure, we performed differential abundance analysis comparing these groups. Since multiple metals co-occur in New Caledonian ultramafic soils and individual metal contributions are difficult to isolate, our approach captures the overall proteome response to metal mixtures, with focused examination of correlations between protein abundance and Ni and Cr concentrations, the primary contaminants of concern in this context.

## 2. Materials and Methods

### 2.1. Sampling Location

New Caledonia is a French island in the South Pacific that hosts approximately 7% of global nickel (Ni) reserves. Nickel mining has occurred on the island since the late nineteenth century. The construction of a smelter in Nouméa, the capital city, in 1910 resulted in long-term atmospheric emissions of Ni and Cr particulates. Previous studies have documented elevated soil metal concentrations across Nouméa, characterised by a spatial gradient in which Ni and Cr concentrations decrease with increasing distance from the smelter [7, 8]. In this study, we selected 18 sites, comprising 17 sites distributed across urban Nouméa along a smelter-related contamination gradient and one site in the mining-impacted town of Thio, which is located adjacent to active laterite strip-mining operations in north-eastern New Caledonia. Figure 1 shows our sampling sites in New Caledonia.

**Figure 1.**
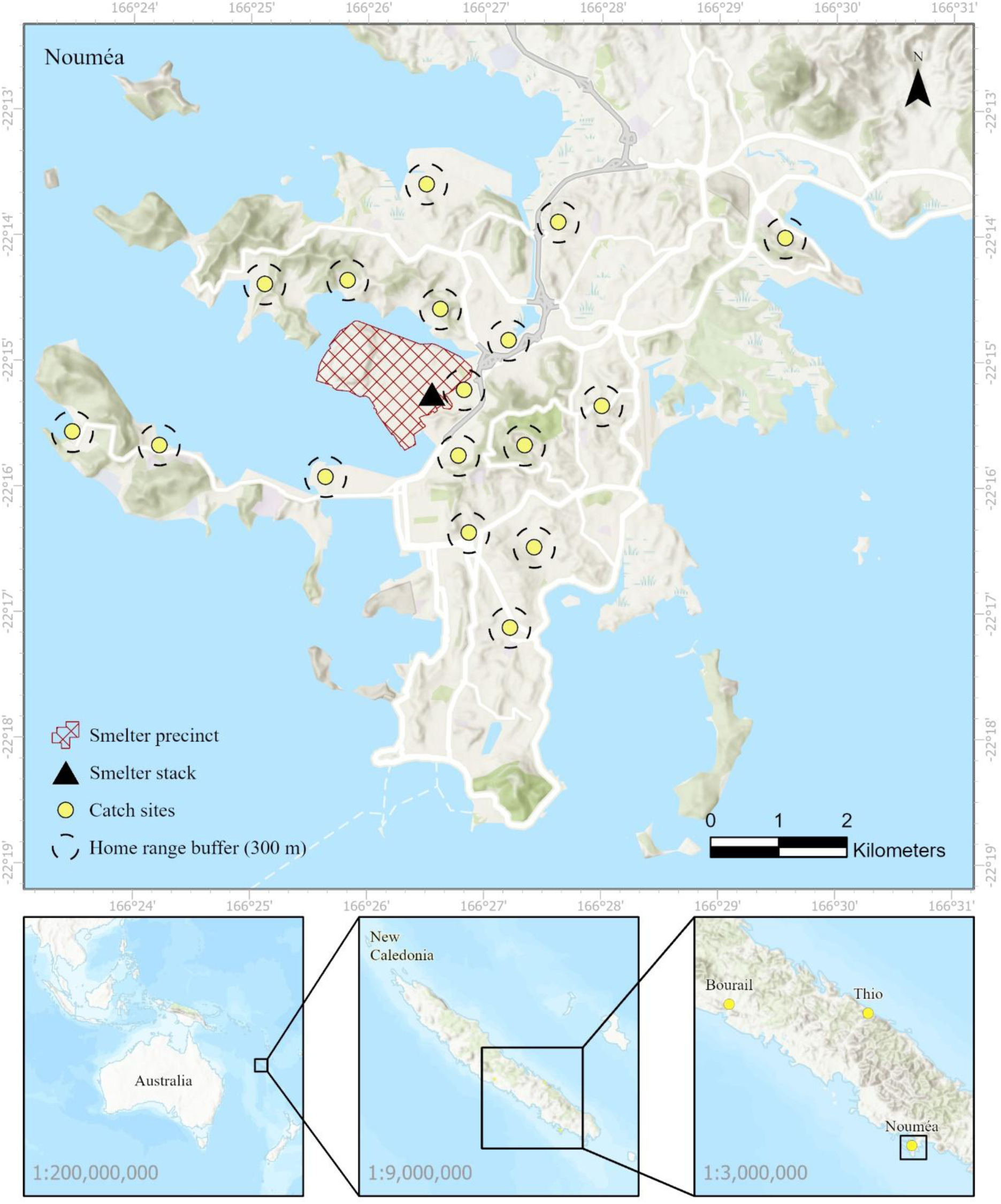
Map of house sparrow capture sites in New Caledonia. Sites within Nouméa were distributed along a contamination gradient with increasing distance from the Ni smelter. The site at Thio represents a mining-impacted location characterised by extreme surface soil Ni and Cr concentrations.

### 2.2. Trace metal measurements

At each site, surface soils (0–2 cm depth) were collected and analysed for metal concentrations following established protocols [21, 25]. Samples were sieved to <250 µm, transferred to Mylar film analysis cups, and analysed for metal concentrations (mg/kg) using X-ray fluorescence (XRF) spectrometry with a Vanta portable XRF spectrometer (Olympus-IMS). We report data on As, Cr, Cu, Fe, Mn, Ni, Pb, Ti, V, and Zn, though we focus on Ni and Cr as the primary contaminants of concern associated with mining and smelting activities in this region. To assess anthropogenic contributions to soil metal concentrations, surface soil values were compared against subsoil background concentrations reported for New Caledonian soils [7]. Ecological Soil Screening Levels (Eco-SSLs) were used to contextualise the potential biological relevance of the observed soil Cr and Ni contamination levels for birds. These screening benchmarks, developed by the U.S. Environmental Protection Agency, are designed to protect sublethal endpoints, such as growth and reproduction, following soil-mediated exposure [26]. Site-specific enrichment factors (EFs) for elements measured in New Caledonian surface soils (2 cm depth). Enrichment factors were calculated using mean element concentrations for each site and normalised to titanium (Ti) as the reference element, following Barbieri (2016). Titanium was selected as a conservative element with minimal enrichment in New Caledonian soils. Enrichment factors represent relative deviation from regional background concentrations [7]. Enrichment factor classification is based on Sutherland 2000 [27].

### 2.3. Biological sampling

Approval for bird capture and sampling was granted by the Macquarie University Animal Ethics Committee (AEC#2020/011-14). At each site, three adult male house sparrows were captured using mist nets. Body mass, wing chord, and tarsus length were measured for each individual. An initial blood sample was obtained by puncturing the brachial vein with a 26-gauge hypodermic needle. Approximately 10 µL of blood was collected using HemoCue Hb 201 Microcuvette Strips and analysed in the field for haemoglobin (Hb) concentration with a HemoCue Hb 201+ point-of-care testing system.

### 2.4. Blood and Plasma collection

House sparrows were euthanized by cervical dislocation, after which approximately 100 µL of blood was collected from the jugular vein. Euthanasia was undertaken to permit the comprehensive collection of multiple tissues required to address several related research objectives. In addition to the samples reported here, tail feathers, red blood cells, plasma, discrete brain regions (cerebellum, optic tectum, and forebrain), and liver tissue (including a small subsample and the remaining liver) were collected from each individual. Blood was transferred into EDTA-treated microcentrifuge tubes and centrifuged at 2,000 g for 10 min. Plasma was then decanted into fresh tubes. Both plasma and cell pellet samples were immediately placed on dry ice and subsequently transferred to a −80 °C freezer for long-term storage. In total, we collected 52 blood samples, of which three plasma samples were lost due to thermal shock, with the remaining plasma samples (n = 49) used in this study. For the pellet, we selected random samples (n = 20) from Nouméa.

### 2.5. Plasma proteome sample preparation

Neat plasma samples were prepared using the method described by Niu *et al.* [28]. Briefly, samples were randomized and defrosted at 4 °C, and 5 µL of plasma was diluted with 45 µL of lysis buffer (10 mM tris(2-carboxyethyl) phosphine, 40 mM chloroacetamide, 100 mM TEAB, pH 8.5) in a 96-well plate. Samples were mixed by pipetting up and down at least 20 times. The plate was centrifuged at 300 *g* for 10 mins at room temperature, and 5 μL of the diluted samples were transferred to a second plate containing 15 μL of lysis buffer. The plate was heated to 95 °C for 10 min using a thermomixer and subsequently cooled for 15 min at room temperature. Trypsin (Trypsin Gold, Promega, Madison, Wisconsin, USA) was diluted to 0.05 μg/μL in 100 mM TEAB (pH 8.5), and 15 μL was added to each well. Samples were centrifuged for 5 min at 200 g and digested at 37 °C for 18 h. Following digestion, peptides were centrifuged at 300 g for 10 min, and 56 μL of 0.2% trifluoroacetic acid (TFA) was added and mixed. Tryptic peptides were desalted using in-house packed stage tips containing SDB-RPS membrane (Empore^TM^ Solid Phase Extraction (SPE) Disks). The SDB-RPS membrane was conditioned in 99% 2-propanol/1% TFA, then equilibrated in 0.2% TFA. Subsequently, 60 μL of tryptic digest was loaded onto the column and the column was washed twice with 0.2% TFA. Peptides were eluted with 80% acetonitrile/5% ammonium hydroxide. Desalted peptides were dried in a vacuum centrifuge and reconstituted in 15 μL of 0.5% formic acid (FA). Peptide concentration was estimated using a NanoDrop spectrophotometer (Thermo Fisher Scientific, CA, USA), and the final concentration was adjusted to 125 ng/μL with 0.5% FA.

Quality control (QC) samples were prepared by pooling 20 randomly selected plasma samples at the beginning of sample preparation. The pooled sample was aliquoted into nine replicates, which were then processed alongside the plasma samples.

### 2.6. Blood cell pellet proteome sample preparation

Blood cell pellet samples were prepared using the optimised Mitra VAMS (Volumetric Absorptive Microsampling) dried blood method described by Molloy *et al.* [29], adapted for a 96-well format. Blood cell pellet samples were randomized and thawed on ice. Blood clots were disrupted by bead beating with glass beads for 20 s. Samples were immediately transferred to ice and centrifuged for 10 min at 5,000 g at 4 °C. Subsequently, 5 μL of the blood pellet was diluted with 100 μL of phosphate-buffered saline (PBS) at a 1:20 ratio (blood pellet: PBS). Diluted blood pellet (35 μL) was transferred to a 96-well U-bottom plate. VAMS devices were removed from specimen bags and placed onto the samples until fully saturated. VAMS tips were allowed to dry at room temperature for 10 min, after which the devices were transferred back to specimen foil ziplock bags containing desiccant and incubated at 4 °C for 4 h. Following incubation, devices were removed from the specimen bags, and the tips were detached from the spindles and placed into a 96-deep-well filter plate (Nunc DeepWell 96-well plate), which was sealed with a sealing mat. Washing buffer (800 μL; 500 mM LiCl, 100 mM Tris) was added through the bottom of the plate wells. The bottom opening was sealed with a second mat, and the tips were washed in a mixer at 300 rpm for 24 h. The bottom mat was then removed, and a deep-well collection plate was placed beneath the filter plate, which was then centrifuged for 4 min at 2,000 g. The tips were further washed three times with 800 μL of washing buffer for 4 min at 700 rpm each, followed by centrifugation at 2,000 g for 4 min after each wash. Following the final wash, dried tips were transferred to a 96-well plate, and 130 μL of lysis buffer (1% sodium deoxycholate, 100 mM triethylammonium bicarbonate, 10 mM tris(2-carboxyethyl) phosphine, 40 mM chloroacetamide) was added. The plate was heated to 95 °C for 10 min. Upon cooling, 1 μg of trypsin (Trypsin Gold, mass spectrometry grade; Promega, Madison, Wisconsin, USA) was added to each well, and samples were digested at 37 °C for 18 h. The trypsin reaction was quenched by adding 200 μL of 0.5% formic acid. Tips were removed from the digest using forceps, and the plate was centrifuged at 2,000 g to pellet sodium deoxycholate. The supernatant was desalted using an Oasis HLB 96-well plate (10 mg sorbent; Waters Milford, MA, USA) as described previously [30].

Quality control (QC) samples were prepared by pooling five randomly selected blood cell pellet samples at the beginning of sample preparation. The pooled sample was aliquoted into three replicates, which were then processed alongside the experimental samples.

### 2.7. LC-MS/MS for single injection

Tryptic peptides (250 ng) from plasma and blood cell pellet samples were analyzed on a Vanquish Neo UHPLC system (Thermo Fisher Scientific, CA, USA) coupled to an Orbitrap Exploris 480 mass spectrometer (Thermo Fisher Scientific, CA, USA). Peptides were loaded onto an Acclaim PepMap 100 C18 trap column (5 μm particle size, 100 Å pore size, 300 μm × 5 mm; Thermo Fisher Scientific, CA, USA) and separated on a 300 mm C18 analytical column (75 μm internal diameter) packed with Dr. Maisch Reprosil-Pur resin (3 μm particle size, 120 Å pore size). Both trap and analytical columns were maintained at 45 °C. Peptides were loaded in buffer A (0.1% formic acid) and eluted using buffer B (80% acetonitrile, 0.1% formic acid) with the following gradient profile: 2% to 37.5% buffer B over 60 min (linear gradient), 37.5% to 95% buffer B over 3 min, and held at 95% buffer B for 10 min. Peptides were electrosprayed using a stainless-steel emitter (Nano Bore O.D. 150 um, I.D. 30 um, Thermo Fisher Scientific, CA, USA). Data were acquired in data-independent acquisition (DIA) mode. MS1 scans were acquired at a resolution of 120,000 with a scan range of 350–1,450 m/z. Maximum injection time was set to automatic with automatic gain control set to standard. Each MS1 scan was followed by 20 MS2 scans using variable overlapping wide windows spanning 350–1,450 m/z. Precursor ions were fragmented using higher-energy collisional dissociation (HCD) at 27% normalized collision energy, and fragment ions were analyzed in the Orbitrap at 30,000 resolution.

### 2.8. LC-MS for multi-injection gas phase fractionation (GPF)

A separate empirical spectral library was generated for each sample type using the same LC-MS system described above. Desalted peptide samples from plasma or blood cell pellet fractions were pooled separately and analyzed using a multi-injection gas-phase fractionation (GPF) strategy. Each pooled sample was injected seven times to cover a precursor m/z range of 395–1,450. The m/z ranges covered by each sequential injection were: 395–505, 495–605, 595– 705, 695–805, 795–905, 895–1,005, and 1,005–1,450. Chromatographic separation conditions were identical to those used for single-injection analysis. MS1 spectra were acquired at 60,000 resolution, and precursor ions were fragmented using higher-energy collisional dissociation (HCD) at 27% normalized collision energy. Automatic gain control was enabled with a normalized AGC target of 1,000 and automatic maximum injection time. Each MS1 scan was followed by 51 staggered MS2 scans. For the first six injections, 4 m/z isolation windows were used, while the final injection (1,005–1,450 m/z) employed 51 isolation windows of 28 m/z to cover the high-mass precursor range. MS2 spectra were acquired at a resolution of 30,000.

### 2.9. Spectral processing

Thermo RAW files were processed using the DIA-NN computational platform (v2.2.0) [31] through a three-step DIA spectral analysis workflow. A previously published high-quality *Passer domesticus* protein database (14,260 sequences) was used for spectral matching [32, 33]. The database contained custom headers and lacked protein and gene annotations. To address this, the database was mapped and annotated using eggNOG-mapper [34], and the headers were subsequently formatted to UniProt-style entries using custom R scripts. When a gene or protein name mapped to multiple sequences, the corresponding annotation was excluded to avoid ambiguity, and the original custom identifier was retained. Consequently, each protein sequence retained a unique identifier, and no two sequences shared the same protein or gene name. Sequences without annotation matches retained their original custom identifiers.

In the first step, this curated FASTA file was used to generate a predicted in silico library using DIA-NN neural networks. MS2 mass accuracies were set to 10 ppm, match-between-runs (MBR) was enabled, and the following parameters were applied for in silico library generation: trypsin as the protease with one missed cleavage allowed; fixed modifications included carbamidomethylation of cysteine and N-terminal methionine excision; no variable modifications were selected to limit the search space; peptide length of 7–30 residues; precursor charge range of 2–4; precursor m/z range of 350–1,450; and fragment m/z range of 200–1,800. In the second step, the in-silico library was used to search multi-injection gas-phase fractionation (GPF) files separately for each sample type. MS1 scan windows were optimized automatically by DIA-NN. Precursors were filtered at 1% false discovery rate (FDR), and proteotypicity was specified using gene names from the database. This process generated sample-specific empirical spectral libraries. In the third step, these empirical libraries were used to search single-injection DIA files with identical parameters, except that MBR was disabled. Quantification was performed using the QuantUMS (high-precision) strategy [35] with cross-run normalization.

### 2.10. Differential abundance analysis

All proteomics data analyses were performed in R (version 4.5.2) using RStudio unless specified otherwise. The plasma ‘pg_matrix.tsv’ output file from DIA-NN was used to check for red blood cell (RBC) and platelet contamination using Baize [36]. Protein differential expression analysis (DEA) for plasma and blood cell pellet samples was performed using the peptide-level quantification model MS-EmpiRe [37] implemented in the MS-DAP R package (version 1.2.2) [38] with the following settings: minimum detections = 2, minimum quantifications = 2, minimum peptides per protein = 2, filter by contrast = TRUE, normalization algorithm = variance stabilizing normalization (VSN) with between-protein mode. MS-EmpiRe was used as the differential expression algorithm with a q-value threshold of 0.05 and a log₂ fold-change cut-off of 0.5. Proteins initially identified as significant were further filtered using the DEqMS model, which is also implemented in MS-DAP. Only proteins with a protein-level p-value < 0.05 from the DEqMS quantification model were considered significant.

MS-DAP also performs differential detection analysis for proteins that lack peptides with sufficient data points across samples for DEA but show substantial differences in the number of detected peptides between sample groups. The following filters were applied for differential detection analysis: a minimum of 2 peptides per protein, a minimum of 4 samples observed, and a minimum fraction observed of 0.5. Proteins with a z-score > 6 or < −6 were considered differentially detected.

### 2.11. Data pre-processing for principal component analysis and dose-dependent analysis

The plasma protein MaxLFQ intensity matrix was generated from the primary ‘report. parquet’ file using the DIA-NN R package with the following quality thresholds: Q.Value ≤ 0.01, PG.Q.Value ≤ 0.05, Global.Q.Value ≤ 0.01, and Global.PG.Q.Value ≤ 0.01. The resulting protein group dataset was filtered in two steps. First, protein groups identified by fewer than two peptides were removed. Second, protein groups with more than 40% missing values across all samples were excluded, retaining only proteins with at least 60% valid quantification values for downstream analysis. Protein intensities were log₂-transformed, and missing values were imputed by random sampling from a normal distribution with a mean downshifted by 1.8 standard deviations (SD) and a width of 0.3 SD relative to the abundance distribution of all proteins within each sample [28]. The blood cell pellet dataset was processed identically. The coefficient of variation (CV) was calculated for each protein in the final matrix before imputation.

### 2.12. Liver sample preparation and analysis

Liver samples were prepared and analysed as described in chapter 3 for the field trip study. Samples were prepared and analysed alongside the samples from Broken Hill and Port Pirie.

### 2.13. Integrated Principal Component Analysis (PCA)

Principal component analysis (PCA) was performed for each dataset using the top 100 most variable features with the mixOmics R package (version 6.34.0) [39]. Confidence ellipses were drawn at 95%, and loadings (proteins) with absolute values > 0.7 were displayed on the plot with arrows representing the direction of loadings driving the PCA.

Samples (*n* = 18) with both plasma and blood cell pellet data available were used for integrated multi-omics analysis. Multi-omics PCA was performed using a concatenated matrix approach. For each omics layer, the top 100 most variable proteins were identified and arranged as feature × sample matrices, which were then transposed to sample × feature orientation. To account for differences in measurement scales between plasma and pellet samples, each matrix was standardized using *z*-score normalization (mean = 0, SD = 1). The two standardized matrices were then horizontally concatenated to create an integrated multi-omics matrix with dimensions *n* × (*p*₁ + *p*₂), where *n* is the number of samples and *p*₁ and *p*₂ represent the number of features from plasma and pellet, respectively. PCA was performed on the integrated multi-omics matrix using the prcomp() function (center = TRUE, scale = FALSE) in R, and plots were generated using ggplot2.

### 2.14. Metal Network Analysis

Soil samples were analyzed using X-ray fluorescence (XRF) spectrometry, and concentrations for 21 elements were obtained for each site. For sites with multiple measurements, element concentrations were averaged. Pairwise associations between metal concentrations were assessed using Spearman’s rank correlation coefficient (*ρ*). Correlations with |*ρ*| > 0.5 were used to construct a weighted, undirected network using the igraph R package. Nodes represented elements, and edge weights corresponded to absolute correlation coefficients. The network was visualized using the Fruchterman-Reingold layout algorithm, and Community detection was performed using the Louvain algorithm to identify groups of co-occurring metals.

### 2.15. Protein - Metal correlation analysis

To identify proteins whose abundance responds to soil metal contamination, Spearman rank correlation analyses were performed between individual protein intensities and soil metal concentrations (Ni and Cr) across all samples. Correlations were calculated using the cor.test() function in R and considered statistically significant if *p* < 0.05 and the correlation coefficient > 0.5. Linear regression models were fitted to quantify dose-response relationships for proteins with significant correlation with Ni and Cr. The linear model was fitted as:

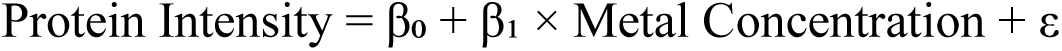

where β₀ is the intercept, β₁ is the slope (dose-response coefficient), and ε represents residual error. Models were fitted using the lm() function in R. Model assumptions (normality of residuals and homoscedasticity) were assessed using Q-Q plots and residual plots. Models violating these assumptions were excluded from further analysis.

### 2.16. Plasma proteome classification, Gene Enrichment, and Network analysis

Gene names mapped via eggNOG-mapper from the final filtered matrices were matched against the Human Protein Atlas plasma proteome database (3,750 proteins). A protein was classified as a plasma protein if its gene name matched an entry in the human database. The absence of a comprehensive avian plasma proteome reference necessitated this approach. Gene Ontology (GO) enrichment analysis was performed on the proteomics dataset using the clusterProfiler package (version 4.18.4) in R, with the *Gallus gallus* reference database (org.Gg.eg.db, version 3.22.0). GO terms for biological process and molecular function categories were assessed, and enrichment terms with adjusted *p*-value < 0.05 (Benjamini–Hochberg correction) were considered statistically significant

## 3. Results

In this study, we aimed to characterize the blood proteome of house sparrows (*Passer domesticus*) inhabiting heavy-metal-contaminated areas in New Caledonia. The island is characterized by elevated levels of nickel (Ni) and chromium (Cr), arising from both natural and anthropogenic sources. Biological samples (plasma, blood pellet, and tissue samples) were collected from adult male house sparrows, and soil samples were collected at the sites of biological sampling. Based on Ni and chromium Cr levels in the soil, samples were divided into three groups: high-contamination group (n = 16), low-contamination group (n = 16), and medium-contamination group (n = 17). We then performed untargeted proteomics analysis on plasma (n = 49) and plasma-depleted blood cell pellets (n = 20), and an overview of the study is shown in Figure 2.

**Figure 2.**
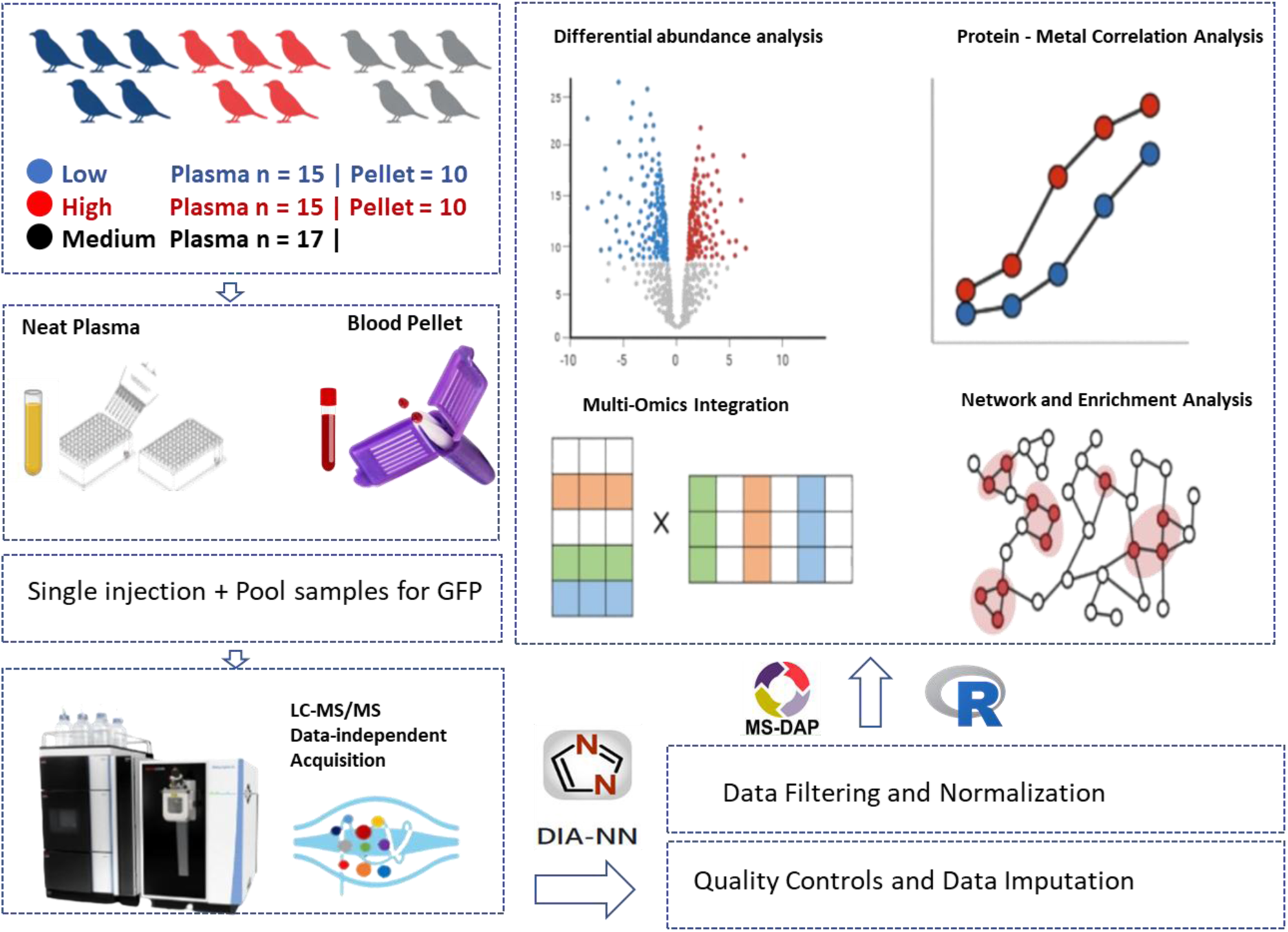
Experimental workflow for house sparrow blood proteomics. Sparrow blood was separated into plasma and cell pellet fractions, with samples divided into three groups. Plasma samples were prepared using a neat plasma approach, while blood cell pellet samples were prepared using a Mitra VAMS-based approach. Tryptic peptides were analysed via data-independent acquisition (DIA) on an Orbitrap Exploris 480 coupled to a Vanquish Neo nanoLC system, and the resulting data were processed with DIA-NN before downstream bioinformatics analysis.

### 3.1. Anthropogenic activities are a major source of Ni and Cr contamination in New Caledonia

Soil samples were collected from 18 sites, which were classified into three contamination levels (low, medium, and high) based on environmental assessments conducted in previous studies [7, 8]. Metal concentrations for each site are provided in Supplementary Tables 1 and 2. Soil Cr concentrations ranged from 402.5 ± 177.3 mg/kg at low contamination sites to 757.4 ± 318.2 mg/kg at medium sites and 2,214.3 ± 3,180.2 mg/kg at high contamination sites. Cr concentrations were significantly higher in medium and high sites compared to low sites (ANOVA: F = 4.26, p = 0.020, Figure 3A). All sites were heavily polluted with Cr relative to both background Cr(III) levels (106 mg/kg) and recommended avian Eco-SSL thresholds for Cr(III) (78 mg/kg) (Figure 3A). Ni concentration followed a similar pattern with significant differences across contamination levels (ANOVA: F = 5.07, p = 0.010, Figure 3B). Mean Ni concentrations increased from 163.4 ± 96.7 mg/kg at low sites to 409.9 ± 179.2 mg/kg at medium sites and 904.0 ± 1,128.2 mg/kg at high sites. Ni concentrations, compared to Cr, were relatively lower and fell below the background (180 mg/kg) and Eco-SSL guidelines (210 mg/kg) at some sites, namely 1, 3, 4, 16, and 18.

**Figure 3.**
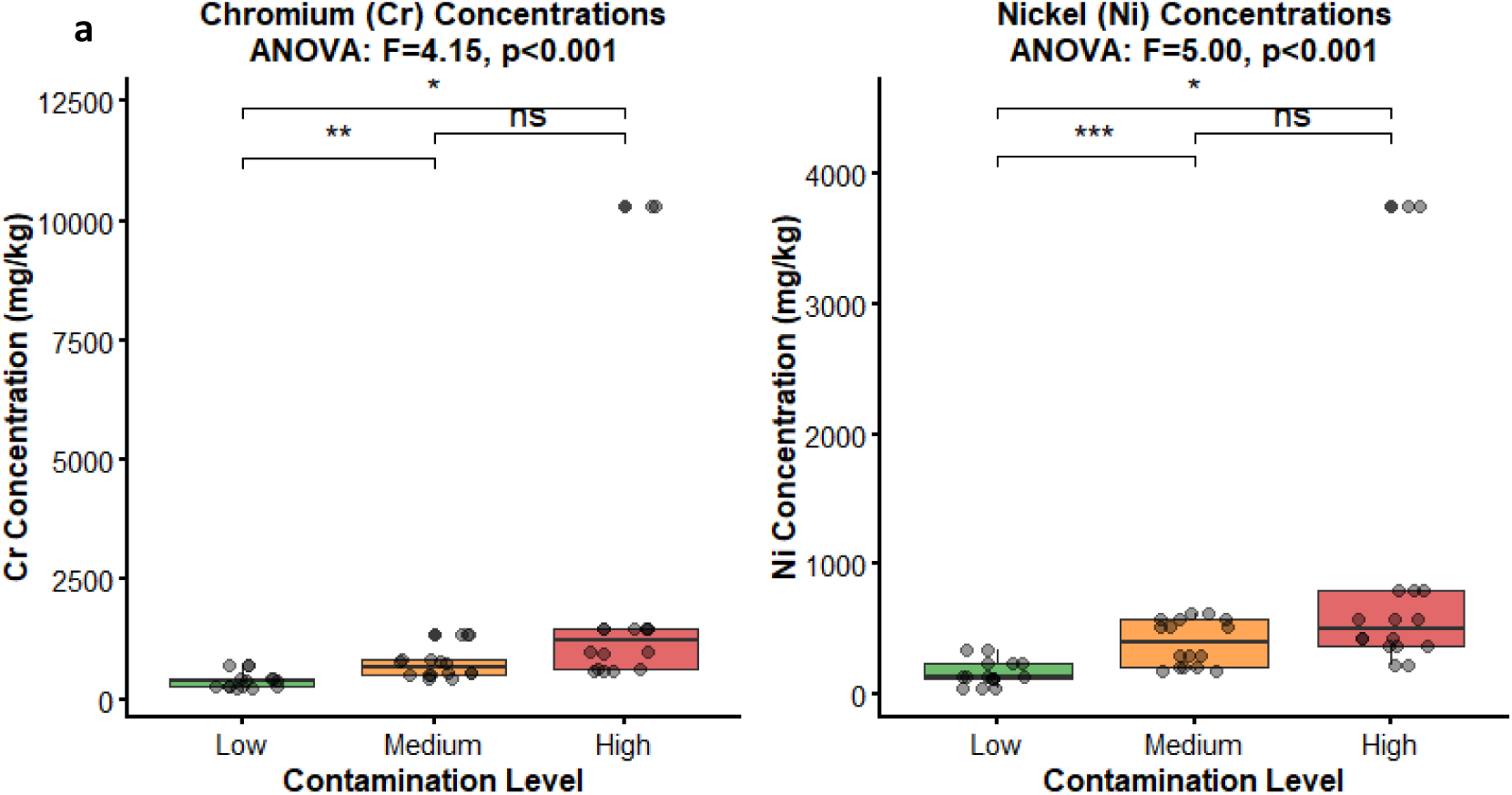
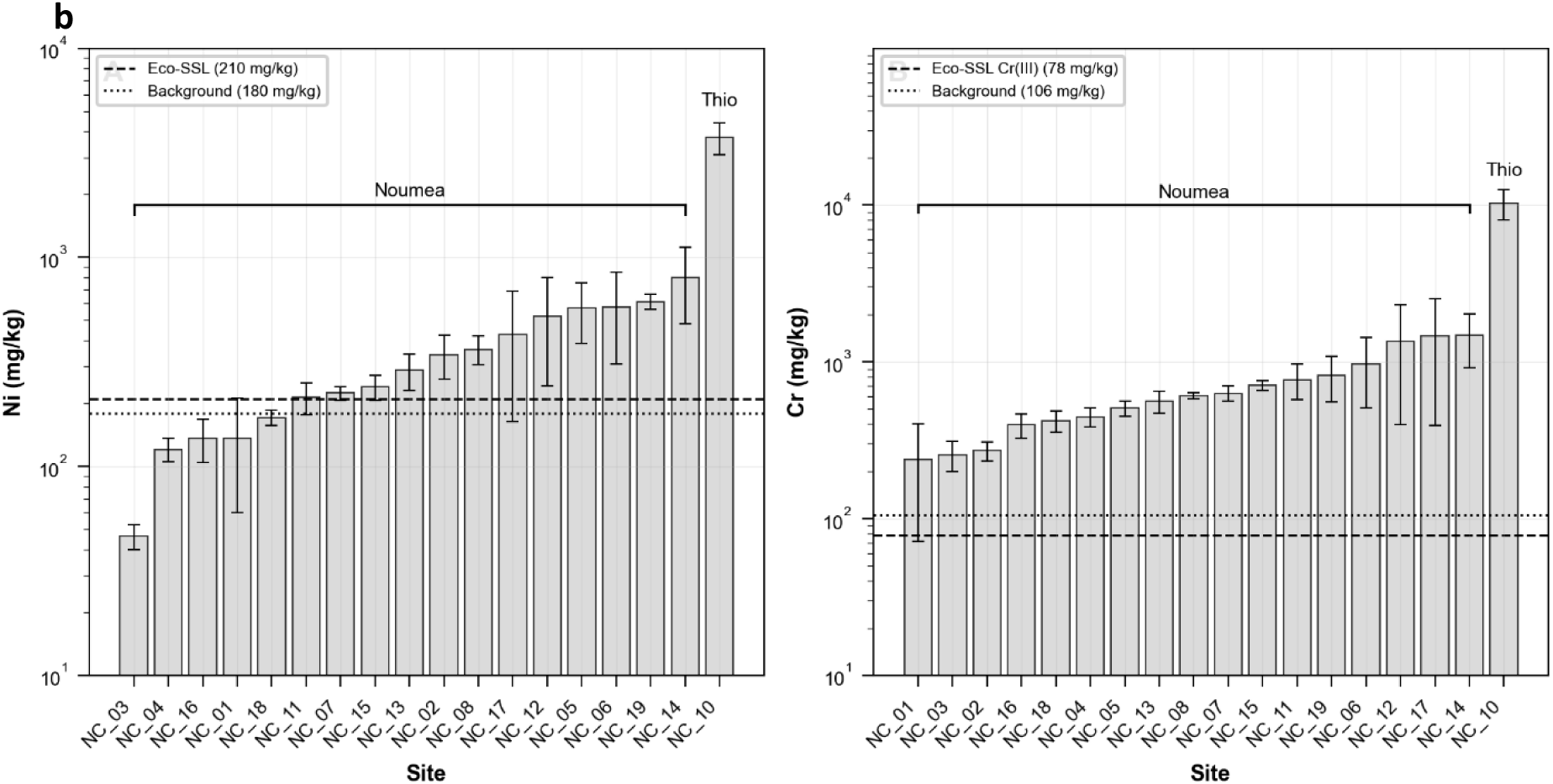
A) Cr and Ni levels in our classification. B) Site-wise levels of Cr and Ni

The Thio mining region was highly contaminated, with mean Cr concentrations 14 times higher than in Nouméa (10,306.7 mg/kg vs. 733 mg/kg) and Ni concentrations 10.5 times higher than in Nouméa (3,754.8 mg/kg vs. 356 mg/kg). There was no significant difference between medium and high contamination levels for both metals (p = 0.069 for Cr and p = 0.084 for Ni). The soil composition indicated that avian species in medium-and high-contamination areas are exposed to significantly higher levels of Ni and Cr than those in low-contamination sites. Since the medium and higher levels did not differ significantly, we anticipated a similar response in the blood proteome.

To identify anthropogenic sources of toxic metals, we performed enrichment factor (EF) analyses for 10 metals at each site, using titanium as the reference element (Supplementary Table 3). Following Sutherland [27], EF values of 2–5 indicate moderate, 5–20 significant, 20–40 very high, and ≥40 extreme anthropogenic enrichment; site means, and individual metal concentrations are shown in Figure 4A. All the sampled sites had EF >2 for Cr and most sites for Ni. As expected, Thio was the most enriched region and showed significantly high enrichment in Ni, Cr, Fe, Zn, and Pb. The EF analysis indicates anthropogenic activities are the major source of Ni, Cr, and other heavy metals. We then performed network analysis of soil metal concentrations to identify which metals cluster together. The analysis identified 5 distinct communities (Figure 4b). Anthropogenic heavy metals (Cr, Ni, Cu, Zn, Pb) formed a tightly interconnected cluster (Groups 1 and 2), indicating a common contamination source. These industrial metals showed positive associations with lithogenic elements (Fe, Mn, Ti, V, As; Group 2, blue) but negative correlations with natural geochemical background elements (Ca, P, S, K, Sr; Group 4, purple), suggesting contamination-related alterations of soil chemistry.

**Figure 4.**
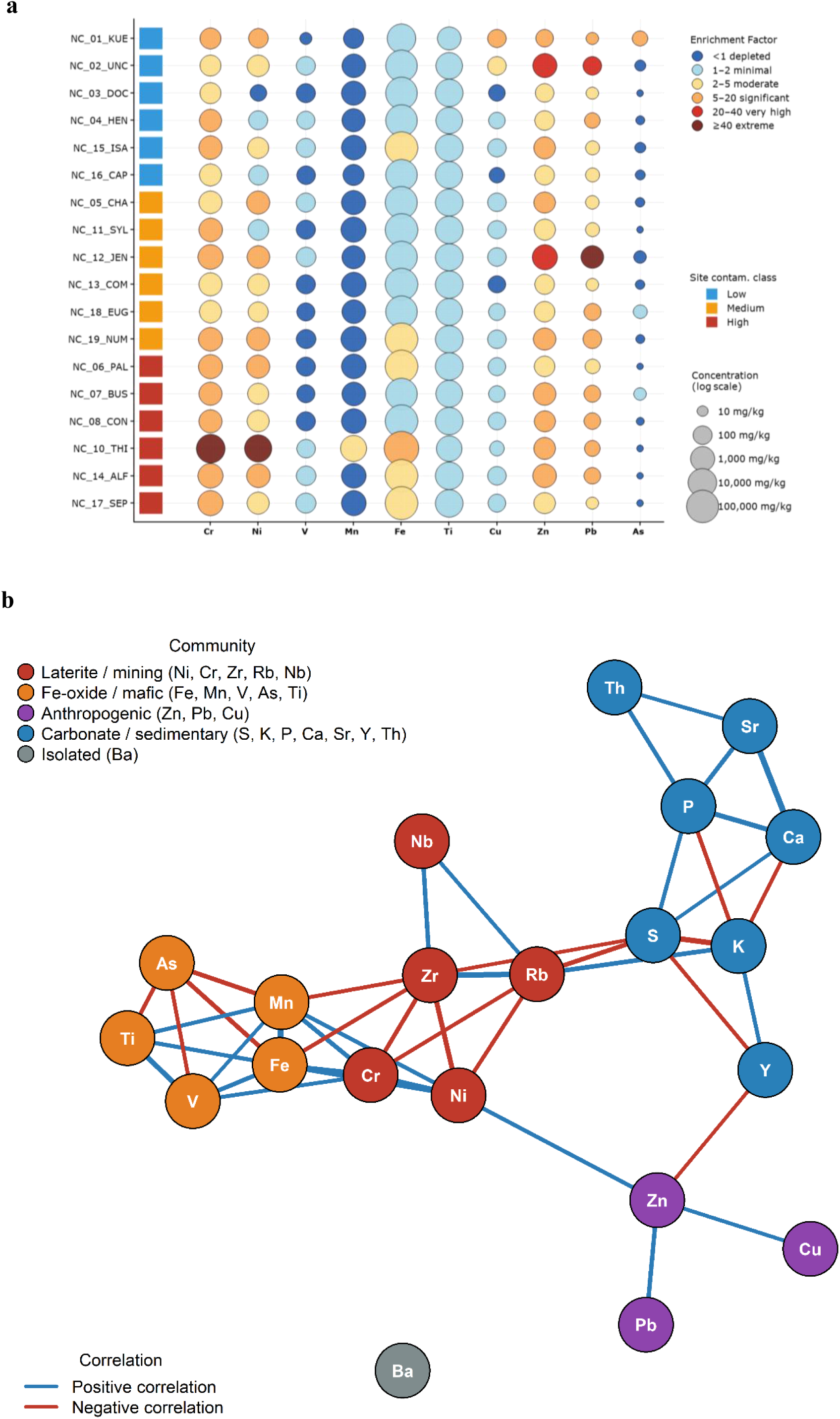
Metal Enrichment Analysis: A) Enrichment factor analysis of heavy metals in New Caledonia. B) Network analysis of soil metal correlations across study sites. Nodes represent individual metals, coloured by community detection, and edge colours indicate correlation direction (blue: positive; red: negative), with edge width representing correlation strength.

### 3.2. Overview of proteomics datasets

No significant differences in biometric measurements (tarsus length, wing length, or body mass) were observed among contamination groups, nor were any significant correlations found between these parameters and Ni or Cr concentrations. Similarly, haem levels showed no significant correlation with either Ni or Cr levels. All biometric data is provided in Supplementary Table 4. Proteomes of plasma and blood cell pellet samples were analyzed by label-free data-independent acquisition (DIA) mass spectrometry. Empirical spectral libraries were constructed for each sample type using gas-phase fractionation (GPF) of pooled tryptic peptides. Library generation identified 660 protein groups in plasma and 3,990 in blood pellets, with 370 shared between the two proteomes.

Single-injection DIA analysis of 49 plasma samples identified 344 protein groups. Proteins identified by a single peptide were excluded, resulting in 277 high-confidence protein groups. The number of protein groups detected per sample is shown in Figure 5A. The dataset was further filtered to remove proteins with >40% missing values across samples. The final filtered dataset comprised 196 protein groups with 95.5% data completeness; 128 protein groups (65.3%) were quantified in all samples without missing values. Quantification reproducibility was assessed using quality control (QC) samples, with a median coefficient of variation (CV) of 15.6% at the protein group level. Of the 196 proteins, 154 (78.7%) exhibited CV <30% (Figures 5B and 5C). DIA analysis of 20 blood cell pellet samples identified 2,193 protein groups. After excluding single-peptide identifications and proteins with >40% missing values, 1,555 protein groups remained in the final dataset with 95.6% data completeness. Median coefficient of variation was 17.5%, with 1,155 protein groups (74.3%) exhibiting a CV <30% (Figure 5D).

**Figure 5.**
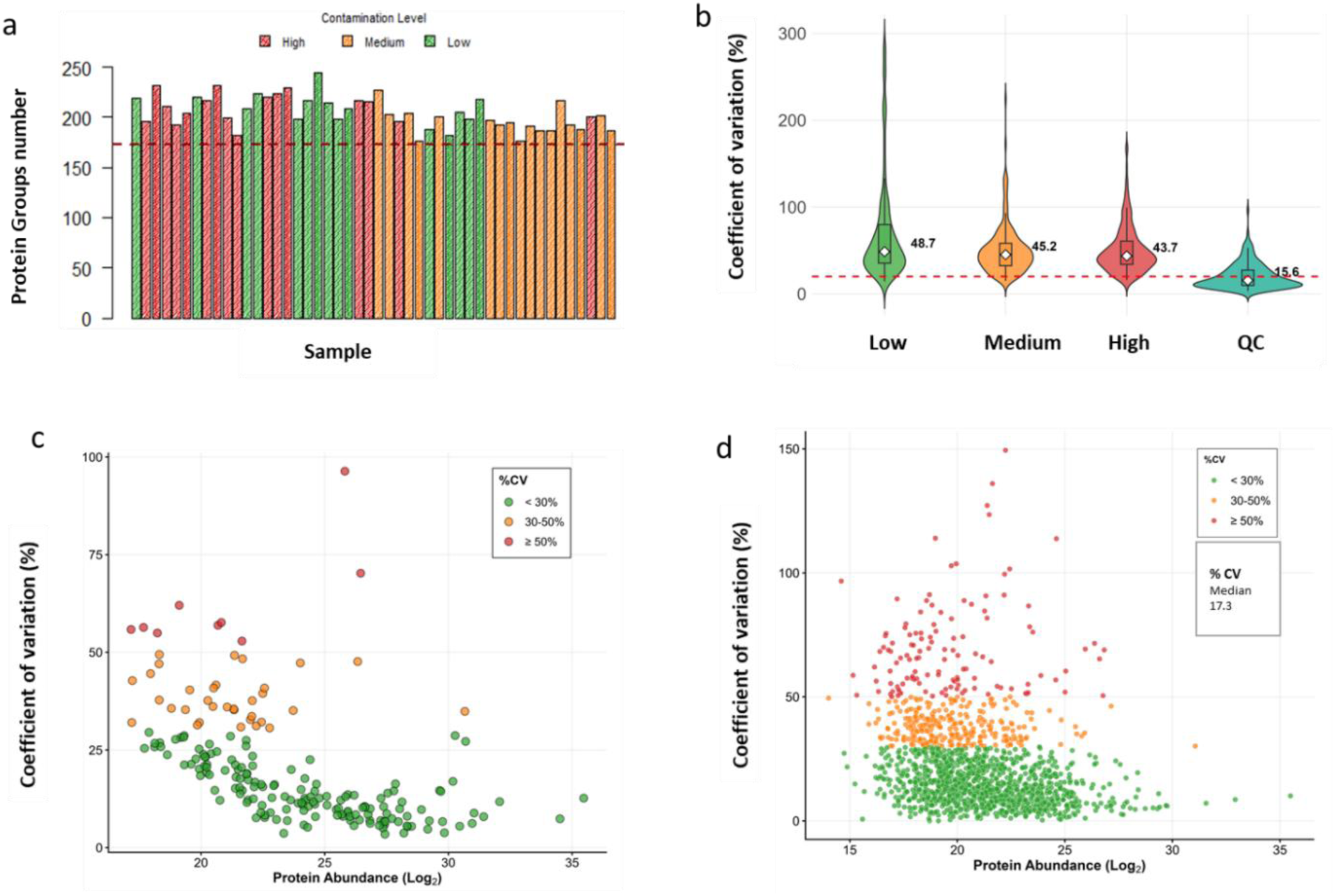
Quality control metrics for plasma and blood pellet proteomic datasets. A) Number of protein groups identified per sample; the red dotted horizontal line indicates the exclusion threshold defined as 2 standard deviations from the median. B) Distribution of coefficient of variation (% CV) across experimental groups from plasma dataset; the red dotted line is drawn at 20%, and the median %CV is printed for each group. C) The relationship between protein abundance (log2 transformed) and % CV in plasma QC samples, each point represents a protein group. Lower-abundance proteins exhibit higher variation due to the high dynamic range of neat plasma. D) same as (c) for blood pellet QC samples. The dynamic range is improved compared to plasma, and % CV is more evenly distributed along the log2 protein intensities.

### 3.3. Pellet depletion strategy enhances detection of plasma proteins in RBC Pellet

A total of 344 protein groups were initially identified in the plasma dataset, of which 188 (55%) were also detected in the pellet fraction. To further investigate the proteomic composition of the pellet samples, subsequent analyses were restricted to confidently identified protein groups, defined as those supported by at least two peptides and present in at least 60% of samples. Applying these criteria retained 196 protein groups in the plasma dataset and 1,555 protein groups in the pellet dataset. We hypothesized that the pellet fraction may contain additional plasma-derived proteins, including proteins not detected in the corresponding neat plasma samples. To test this hypothesis, gene names from both datasets were matched against the Human Plasma Proteome catalogue from the Human Protein Atlas, comprising 3,750 annotated plasma protein genes, to identify proteins classified as bona fide human plasma proteins.

In the plasma dataset, 144 genes (73.5%) matched the database. The lower matching rate may reflect discrepancies between gene nomenclature in eggNOG-mapper annotations and the Human Protein Atlas database, since eggNOG-mapper includes gene names from a diverse range of species. For pellet samples, 694 of 1,548 protein groups (45%) were identified as plasma proteins (Figure 6A). This indicates that blood cell pellets contain a substantial proportion of circulating proteins. Given potential gene-name mismatches, the actual proportion is likely higher. Gene Ontology enrichment analysis indicated that protein localization in both plasma and pellet is highly similar, with the highest enrichment for the extracellular region, extracellular space, and extracellular vesicles (Figure 6B and 6C). To identify RBC-derived proteins, we matched our pellet dataset with a high-quality RBC protein reference from Sea-Lee et al. [40], containing 1202 proteins. Of the proteins detected in our pellet sample, 506 (32.5%) were identified as RBC proteins. Among these RBC proteins, 292 (57.7% of RBC proteins, or 18.7% of total pellet proteins) were also classified as plasma proteins based on the Human Protein Atlas blood proteome database, while 214 (42.3% of RBC proteins, or 13.7% of total pellet proteins) were RBC-specific. The remaining proteins (∼ 42%) may originate from Peripheral Blood Mononuclear Cells (PBMCs). Plasma proteins detected in the pellet were highly correlated with their neat plasma counterparts: of the 86 proteins shared between the two datasets, 17 showed significant correlation (Pearson r > 0.4, p < 0.05, for 18 paired samples), with 16 positively correlated (Supplementary Table 5). This concordance validates the pellet as a reliable source of plasma protein information and confirms that the abundance changes observed in plasma are reflected in the cellular compartment

**Figure 6.**
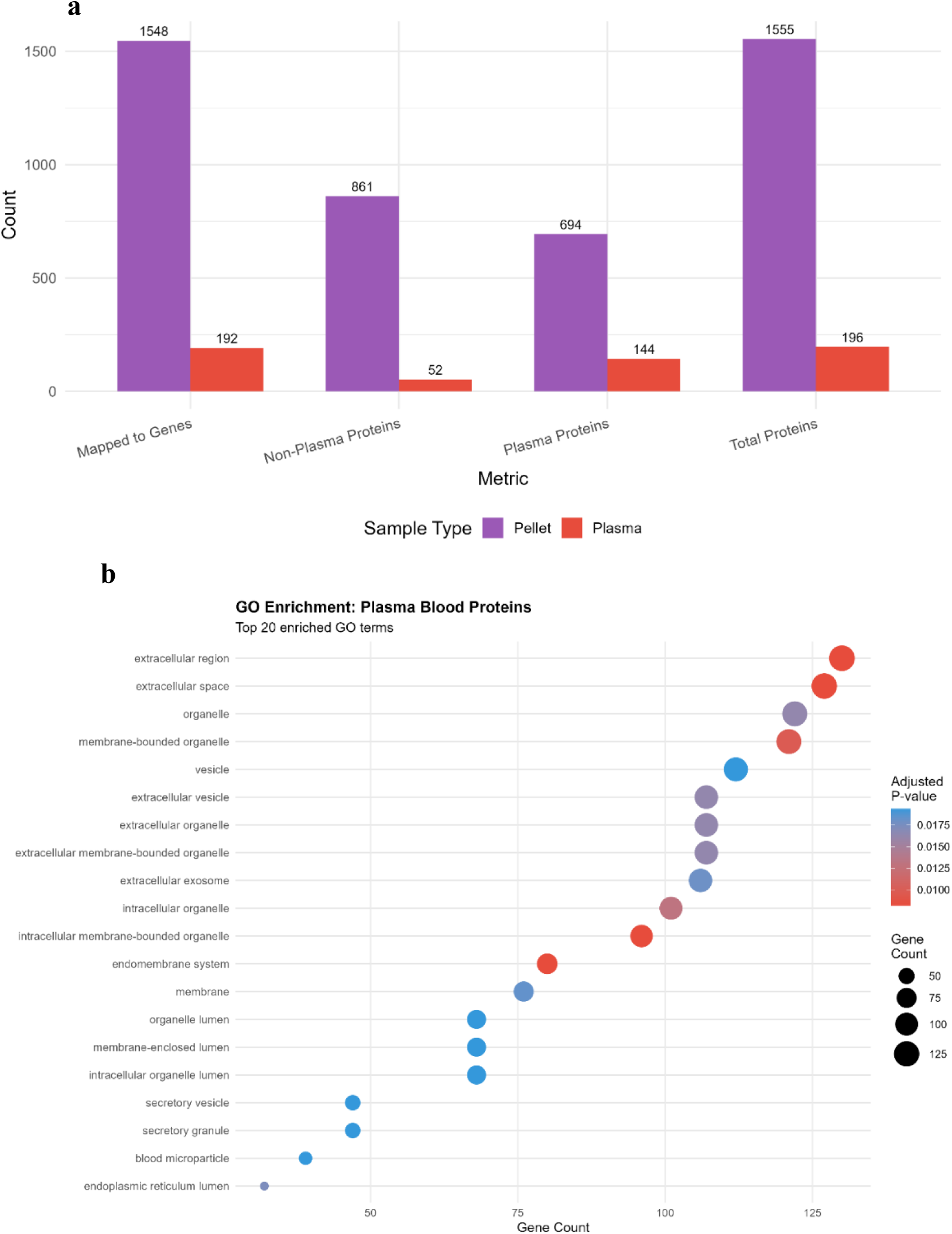
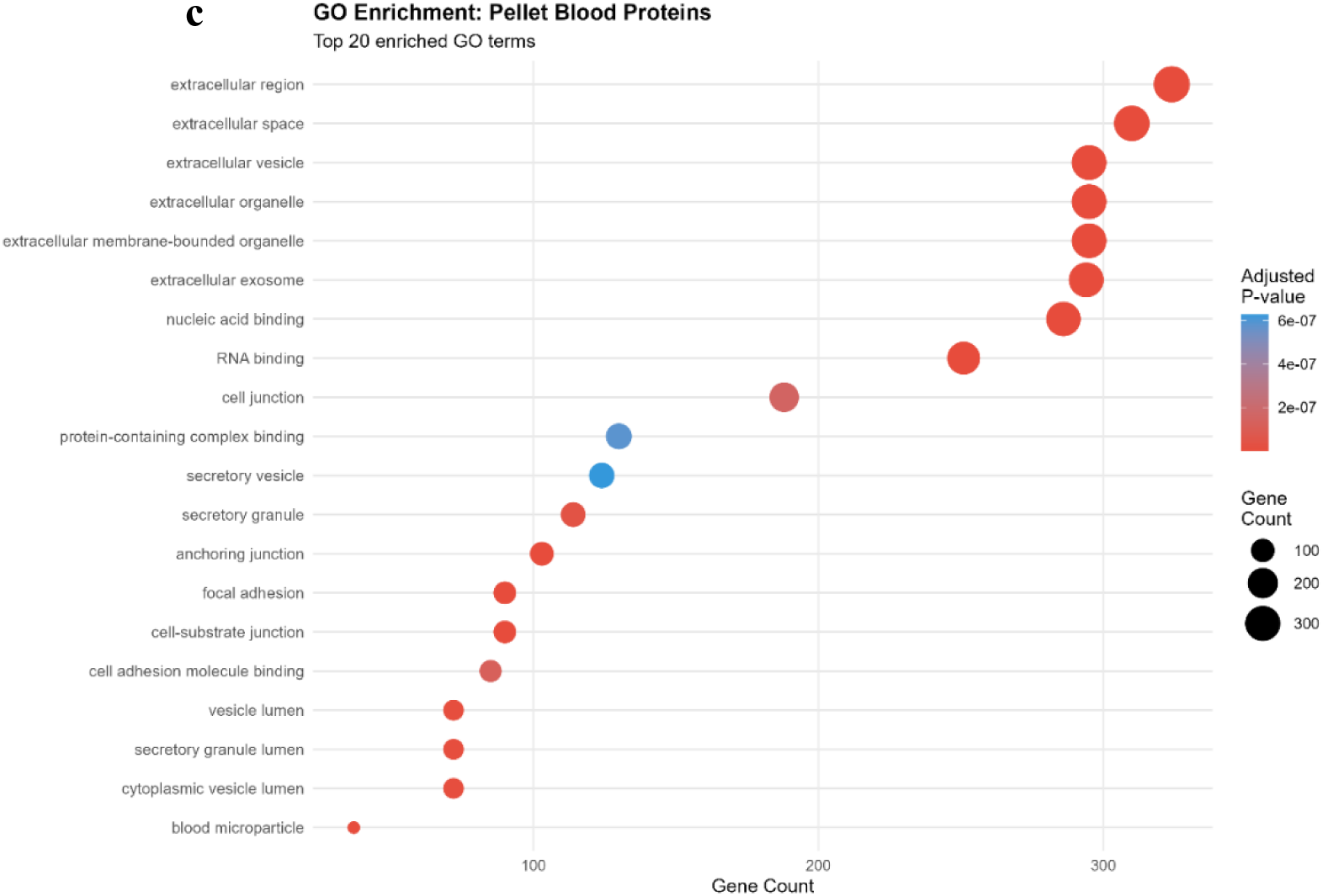
A) Protein classifications as plasma proteins based on Human Plasma Proteome from Human Protein Atlas. Gene Ontology analysis of cellular components for B) plasma and C) blood cell pellet. The significance criterion was an FDR-adjusted p-value < 0.05 (Benjamini–Hochberg correction).

The pellet fraction yielded greater coverage of plasma proteins than neat plasma itself, likely due to reduced interference from high-abundance plasma proteins that typically mask low-abundance proteins in neat plasma. This indicates that depleted pellet samples can provide access to low-abundance plasma proteins that are difficult to detect in neat plasma. Furthermore, the pellet contained proteins from immune cells and other blood cell types that are integral to host responses against pathogens and heavy metals, representing cellular proteome information not captured in cell-free plasma analyses.

### 3.4. Complementary proteomic profiles of plasma and cell pellet enhance biomarker classification through integrated analysis

Principal component analysis (PCA) of plasma and pellet are shown in Figure 7. For the plasma, the medium and high exposed groups separate on the first component (Figure 5A), loadings on biplot showed proteins involved in metabolic process driving the PC1 (Figure 7B) and GO analysis of proteins driving PC1 (cut off >0.15) indicated enrichment in carboxylic acid and oxoacid metabolic processes. Pellet samples were partially separated on PC2 (Figure 7C and 5D). The only enrichment term was serine-type endopeptidase activity. We then integrated samples (n = 18) for which both datasets were available to see whether this improved the overall PCA separation. Groups were visibly more distinct on PC2 (Figure 7E); we then assessed group separation using two metrics: PERMANOVA R² and a separation ratio metric (see methods). Overall group separation improved on both matrices, with PERMANOVA R² (0.217, p < 0.001) and separation ratio 0.746 (Figure 7G). The variables from the pellet were mostly DNA binding proteins, including histone proteins, and in plasma were mainly metabolic processes (Figure 7F). Since Ni is known to induce epigenetic changes by modifying histone proteins, the changes in the abundance of histone and other DNA-binding proteins may be a hallmark of Ni toxicity within the cell [10, 41, 42].

**Figure 7.**
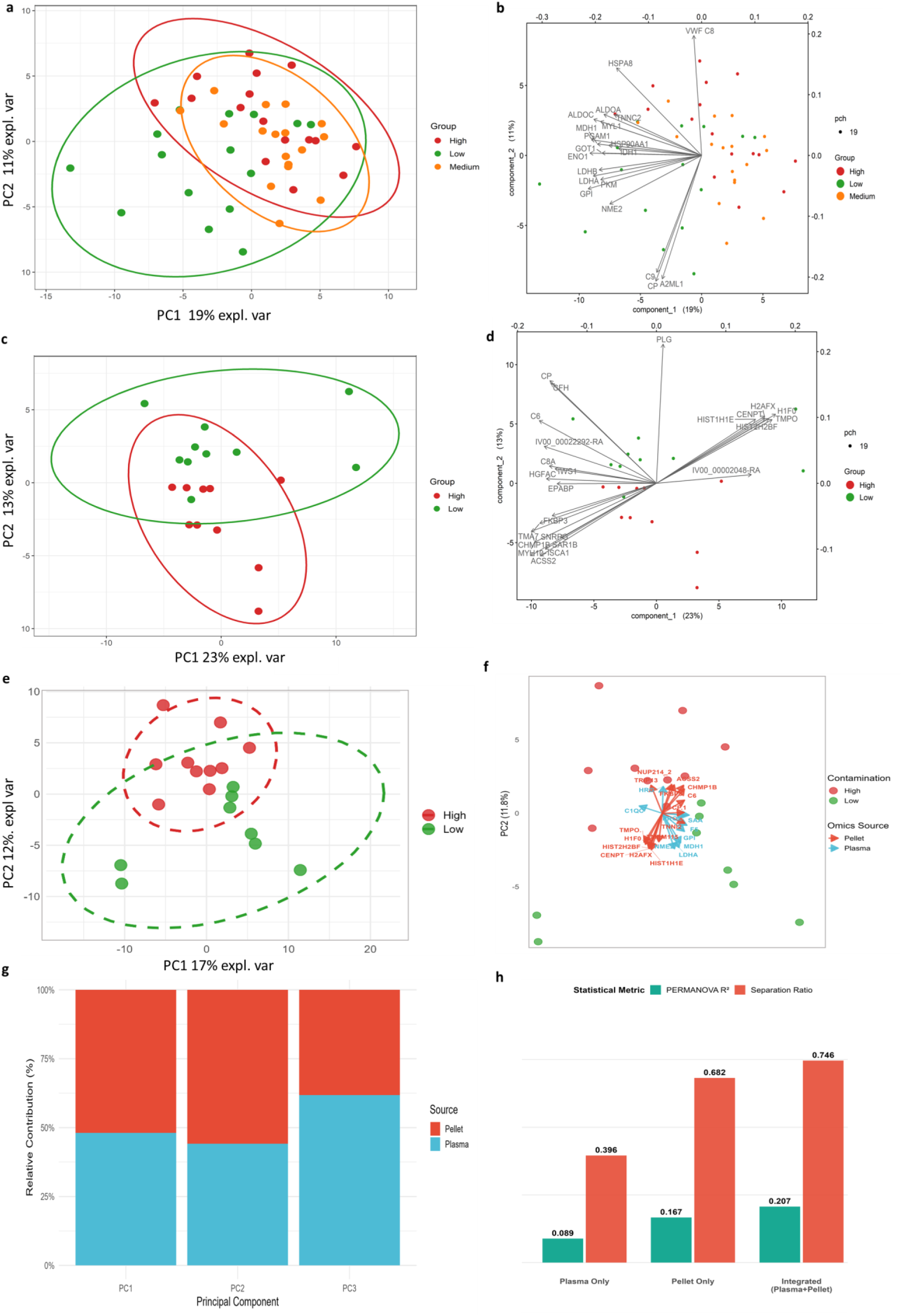
Principal component analysis (PCA) of plasma and blood cell pellet proteomes. PCA was performed on the top 100 most variable proteins from each dataset. Ellipses represent 95% confidence. A) PCA of plasma samples showing separation along PC1 by contamination exposure group. B) Biplot of plasma PCA showing protein loadings contributing most to PC separation. Arrow direction indicates direction of protein abundance increase C) PCA of blood cell pellet samples. D) Biplot of pellet PCA. E) Integrated PCA combining plasma and pellet proteomes from matched samples (n = 18). F) Biplot of integrated PCA. G) Relative contribution of plasma and pellet features to each principal component. H) Statistical validation of multi-omics integration: PERMANOVA R² and separation ratio both demonstrate that integration improves discrimination of contamination groups compared to individual datasets.

### 3.5. Impact of Ni and Cr toxicity on plasma proteome

Pairwise differential abundance analysis was performed using the MS-EmpiRe model in the MS-DAP platform. For pairwise comparisons, 134 proteins were compared between the low-and high-contamination groups, and 126 were compared between the low-and medium-contamination groups. Samples from the high contamination group exhibited a greater extent of proteome dysregulation, with 26 proteins (19.4%) significantly differentially abundant, 4 proteins were increased in abundance, and 22 were reduced in abundance in samples from high contamination areas (Figure 8A). For comparison between low and medium contamination groups, 13 proteins (10.3%) were significantly altered, of which 3 were increased in abundance in the medium contamination group, and 10 were decreased (Figure 8B). Across both comparisons, 11 proteins were consistently dysregulated. This included two proteins: histidine-rich glycoprotein (HRG) and the C-chain subunit of C1q (C1qC), which were more abundant in the heavy metal-exposed groups.

**Figure 8.**
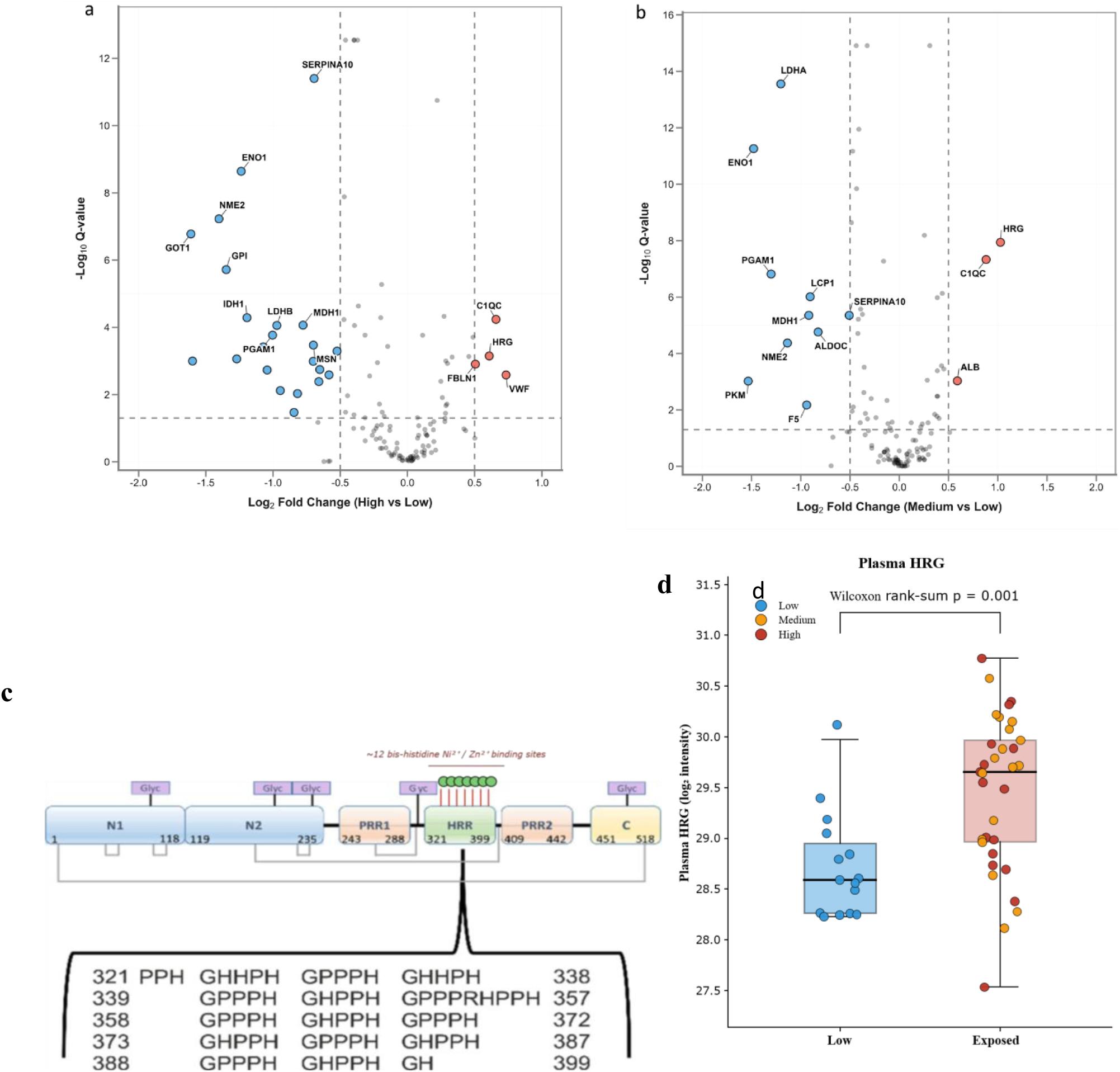
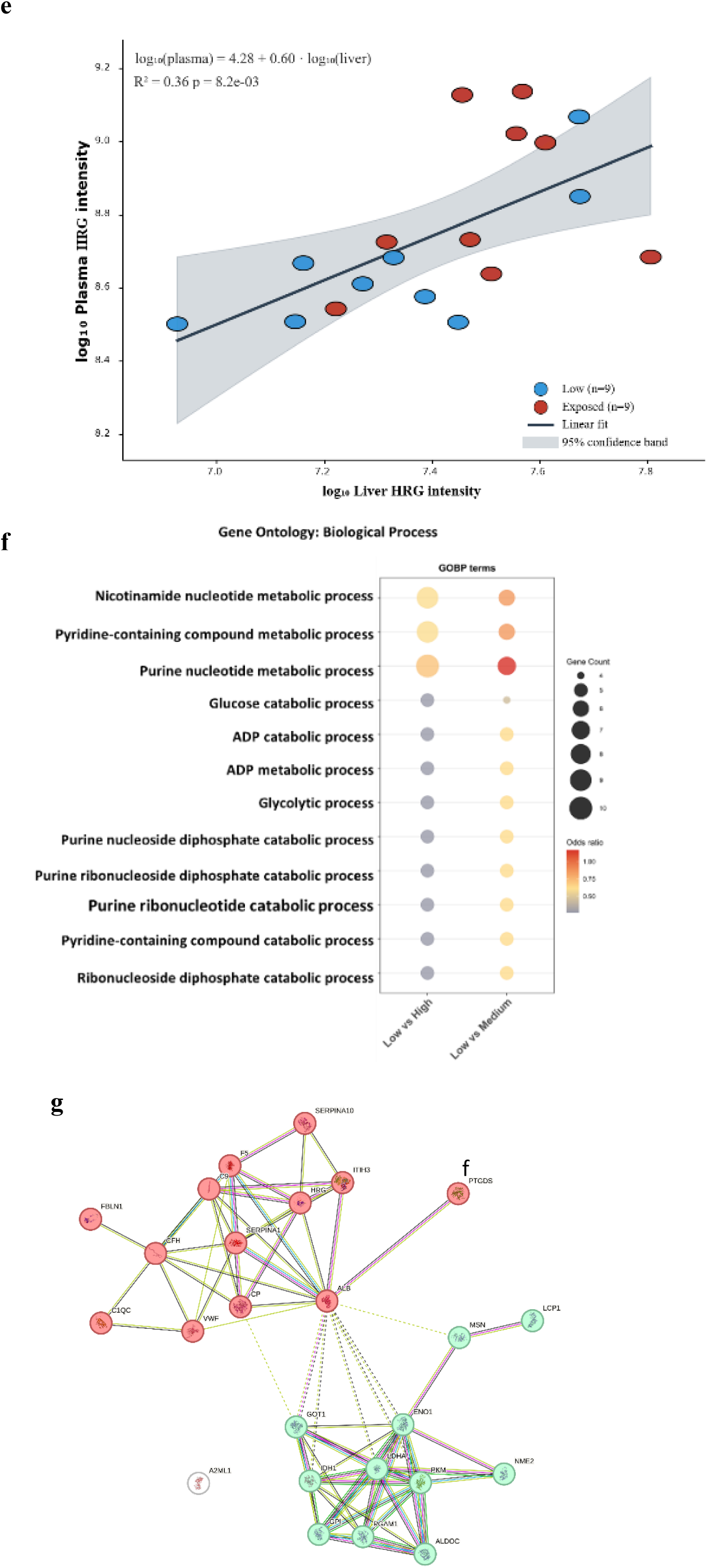
Differential abundance analysis of plasma proteomes across contamination groups. A protein was considered differentially abundant if it met dual criteria: q-value < 0.05 and |log₂ foldchange| > 0.5 in the MS-EmpiRe model, followed by p-value < 0.05 in the DEqMS model. In volcano plots A) and B) the x-axis represents log₂ foldchange, with positive values indicating higher abundance in the contamination group and negative values indicating lower abundance. The y-axis represents −log₁₀(q-value) from MS-EmpiRe, with higher values indicating greater statistical significance. Red points denote significantly upregulated proteins (more abundant in high/medium contamination groups), blue points denote significantly downregulated proteins (less abundant in high/medium contamination groups), and grey points represent non-significant proteins. All comparisons are relative to the low-contamination group as the reference. A) Volcano plot comparing high-and low-contamination groups. B) Volcano plot comparing medium-and low-contamination groups. C) HRG domain architecture, showing the histidine-rich region (HRR) that mediates divalent metal binding. D) Plasma histidine-rich glycoprotein (HRG) levels were compared between the Low and Exposed groups. E) Linear model relating log₁₀ plasma HRG intensity to log₁₀ liver HRG intensity across 18 paired birds. F) Gene Ontology (GO) enrichment analysis showing significantly enriched biological process terms among differentially abundant proteins. G) Protein–protein interaction network generated from the STRING database, revealing two distinct subnetworks that are dysregulated across contamination groups.

HRG is a multidomain plasma protein that binds a wide range of ligands, including divalent metal ions such as Zn²⁺ and Ni²⁺ (Figure 8C). This indicates that the rise in HRG is a direct response to Nickel and even at low levels HRG rises in the plasma (Figure 8D). HRG is primarily synthesised in the liver and secreted into the bloodstream, so the rise in plasma HRG observed in exposed birds most likely reflects an increased synthesis of hepatic HRG production in response to heavy metal load. To test this, we examined HRG in a parallel liver proteomics dataset and performed a paired plasma–liver correlation across 18 birds in which both compartments were quantified. Plasma and liver HRG levels were strongly positively correlated (Pearson r = 0.60, p = 0.008; Spearman ρ = 0.70, p = 0.001), and a log–log linear model gave log₁₀(plasma HRG) = 4.28 + 0.60·log₁₀(liver HRG), R² = 0.36, p = 0.008 (Figure 8E).

In plasma, nine proteins showed consistently lower abundance (LCP1, SERPINA10, MDH1, F5, PGAM1, ENO1, LDHA, NME2, and PKM). Differentially abundant proteins are listed in Supplementary Table 6.

Von Willebrand factor (VWF) and fibulin-1 (FBLN1) were only identified as significantly increased in the high contamination group, while serum albumin (ALB) was only identified as significantly increased in the medium contamination group. The proteins reduced in abundance include serine protease inhibitor proteins (ITIH3, SERPINA1, SPINK5, and A2ML1), complement system proteins (CFH and C9), lipocalin-type prostaglandin D synthase (PTGDS), and ceruloplasmin (CP). Gene enrichment analysis of all significant proteins from each comparison showed very similar enrichment patterns (Figure 8F). The most strongly enriched biological processes were involved in NAD^+^ synthesis and processing. This may indicate a decrease in redox reactions or in the secretion of these proteins by cells, thereby compensating for cellular energy and redox processes, as a consequence of oxidative stress caused by heavy metals. Protein network analysis revealed two interconnected networks (Figure 8G), involving metabolic processes and the immune system/coagulation network [15].

### 3.6. Impact of Ni and Cr toxicity on pellet proteome

Differential expression analysis (DEP) of cell pellet samples between the low contamination group (n = 10) and the high contamination group (n =10) identified 23 proteins that were significantly differentially abundant. Of these differentially abundant proteins, 4 were increased in abundance in the high contamination group, and 19 proteins were decreased in abundance (Figure 9A and Supplementary table 7). Proteins increased in abundance included the C-chain subunit of C1q (C1qC), calcium uniporter regulatory subunit (MCUB), C-1-tetrahydrofolate synthase, cytoplasmic (MTHFD1), and E3 ubiquitin-protein ligase (MYCBP2). Proteins decreased in abundance included histone proteins (H3F3A, HIST2H2BF, H2AFJ, H2AFV), chromatin regulators (HEMGN, CENPT, RCC1, NUCKS1), mRNA splicing processing proteins (DDX17, DHX15, ALYREF), ceruloplasmin (CP), and NF-κB signalling pathway activator SQSTM1 (also known as p62).

**Figure 9.**
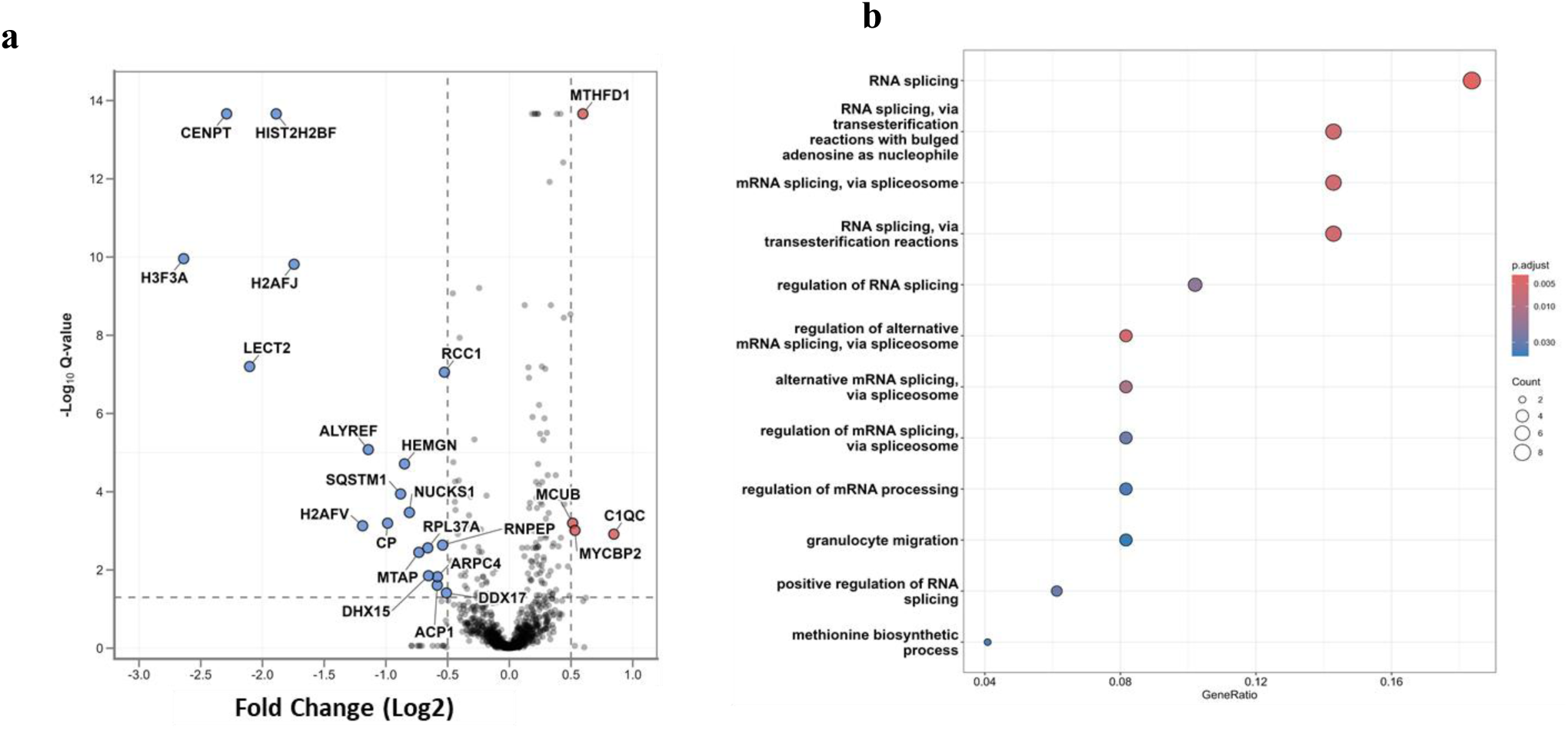
Differential abundance analysis of blood cell pellet proteomes. A) Volcano plot showing differentially abundant proteins between high-and low-contamination groups. B) Gene Ontology Biological Process enrichment from differential abundance and

Differential detection analysis (absolute Z-score ≥6) identified 28 proteins differentially detected between samples, of which 11 were exclusively detected in the high-contamination group (Supplementary Table 8). These included metalloreductase (STEAP3), pachytene checkpoint protein 2 (TRIP13), von Willebrand factor (VWF), and heat shock 70-kDa protein 14 (HSP70). On the other hand, 17 proteins were exclusively identified in the low-contamination group; these included the major acute-phase reactant (SAA1), histone H1.4 (HIST1H1E), and SERPINB10. Gene enrichment analysis using significantly altered genes from DEP and differential detection analysis was enriched for RNA splicing, the most enriched biological process term (Figure 9B), while for molecular function, structural constituent of chromatin and oxidoreductase activity acting on metal ions were the two most enriched terms.

### 3.7. Dose-dependent analysis

Linear models were fitted to identify proteins whose intensities were dependent on soil Ni or Cr levels. We first performed Spearman correlation analysis between protein intensities (log2) and soil Ni and Cr levels (mg/kg). Significant correlations (p-value <0.05 and ρ > 0.5) were then used to fit linear models. For the plasma dataset, no protein passed the significant criteria. From pellet samples, 50 proteins (Supplementary Table 9) were found to have significant correlation with Cr, and 44 with Ni (Supplementary Table 10). Many of these proteins were common to both metals, which may be explained by the strongly correlated concentrations of Ni and Cr in the soil (r = 0.97, p-value = 0.001). It is also worth noting that Fe levels in soil are strongly correlated with Ni and Cr levels (r > 0.95, p-value = 0.001), and the dose-dependent response may be attributable to Fe contamination.

From linear models, a protein was considered significantly dose-dependent if R2 > 0.3 and p-value < 0.05. A total of 21 proteins were significantly dose-dependent for Cr (Supplementary Table 11), and 13 proteins were dose-dependent for Ni (Supplementary Table 12). Of these, 8 were common to both (RNPS1, RBMX, RCC1, ESD, VRK1, HIST1H1E, TRIP13, and RPL37A). Many of the dose-dependent proteins were already identified as differentially abundant or detected, including SERPINB10, RBMX, ALYREF, HIST1H1E, RCC1, HEMGN, RPL37A, TRIP13, and VWF. On the other hand, numerous proteins were only identified through dose-dependent analysis, including VRK1, RNPS1, NUP153, MAN2B1, PFN1, FMNL1, and LSm4.

The dose-dependent analysis indicates that protein dysregulation was associated with the cytoskeleton, DNA damage repair, chromatin remodeling, mRNA processing, and cell cycle control. Cytoskeleton protein abundances decrease with Cr levels in soil, including actin-related protein 2 (ACTR2), actin-binding proteins, Profilin-1 (PFN1), and actin regulatory protein formin-like 1 (FMNL1). Positive correlation with soil Cr levels was also observed for Vaccinia-Related Kinase 1 (VRK1) and Thyroid Hormone Receptor Interactor 13 (TRIP13). In contrast, a negative correlation with soil Cr levels was observed for mRNA-processing proteins (RNPS1, RBMX, ALYREF, LSM4, and RPL37A) and nuclear proteins (HIST1H1E, RCC1, NUP153).

## 4. Discussion

This study investigated blood proteome changes associated with nickel (Ni) and chromium (Cr) exposure in house sparrows (*Passer domesticus*) from New Caledonia using LC-MS-based proteomics of both neat plasma and blood cell pellet fractions. Blood proteomics studies have focused almost exclusively on plasma, overlooking the cellular fraction despite blood cells harbouring critical pathophysiological information. A major barrier has been the overwhelming abundance of haemoglobin and other highly expressed proteins in the pellet. We optimised a volumetric absorptive microsampling (VAMS) based depletion approach enabling deep blood cell proteomics from minimal volumes, revealing that the pellet is substantially enriched in plasma proteins: as a conservative estimate, we quantified 3.5-fold more plasma proteins in the pellet than in neat plasma. This strategy thus provides simultaneous access to proteins originating from plasma, red blood cells, peripheral blood mononuclear cells, and other tissues from a single microsampled blood spot.

Profiling of 49 plasma samples identified 344 proteins, with both exposed groups showing a similar pattern of dysregulation relative to the low-exposure group. Five proteins were significantly increased in abundance: histidine-rich glycoprotein (HRG), complement C1q C-chain (C1qC), albumin, von Willebrand factor (VWF), and fibulin-1, with HRG and C1qC reaching significance in both medium-versus-low and high-versus-low comparisons. HRG and albumin are liver-derived plasma proteins with well-characterised metal-binding capacity [43, 44]. The histidine-rich region of HRG is highly conserved and binds divalent cations, including Zn²⁺, Cd²⁺, Co²⁺, Cu²⁺, and Ni²⁺ [45], while albumin is one of the primary extracellular metal transporters [46]. Their coordinated increase likely reflects enhanced extracellular metal buffering, thereby reducing the bioavailable pool of free Ni and Cr ions. C1qC is a subunit of the C1q complex that initiates the classical complement pathway, and C1q-domain-containing proteins have previously been reported to be upregulated under Cu²⁺ and Cd²⁺ exposure [47]. HRG interacts with C1q to regulate innate immunity and clearance of necrotic cells, raising the possibility that the co-upregulation of HRG and C1qC serves dual roles in metal scavenging and immune response.

In contrast to the upregulated metal-binding proteins, a substantial proportion of liver-derived proteins decreased in the plasma of exposed birds. These included protease inhibitors (ITIH3, SERPINA1), complement regulators (CFH, C9), and the anti-inflammatory proteins prostaglandin D synthase (PTGDS) and ceruloplasmin (CP). Serum amyloid A1 (SAA1), a major positive acute-phase protein that can increase up to 1000-fold during inflammation [48], was also reduced in the blood cell pellet of exposed birds, further reinforcing the pattern of a suppressed rather than activated acute-phase response. The broad decline in liver-origin secretory proteins suggests that chronic exposure to Ni and Cr compromises hepatic protein synthesis or accelerates protein turnover through metal-induced misfolding and degradation.

To investigate intracellular mechanisms of toxicity, we turned to the proteome of the blood cell pellet. Ceruloplasmin (CP), the principal copper-carrying ferroxidase in plasma, was significantly decreased in both datasets. CP catalyses the oxidation of toxic ferrous iron (Fe²⁺) to ferric iron (Fe³⁺), a reaction essential for iron loading onto transferrin and for safe iron efflux from cells [49, 50]. Reduced CP activity has been implicated in pathological iron accumulation in liver, pancreas, and brain tissue [51, 52] and in iron-driven oxidative stress through Fenton chemistry [53, 54]. Elevated levels of Fe in New Caledonia, along with Ni and Cr, may contribute to decreased CP-mediated ferroxidase activity. This reduction would exacerbate environmental iron accumulation and compromise endogenous iron regulation, thereby imposing a dual burden of iron toxicity. The mechanistic basis for CP decline likely involves nickel interference with copper metabolism: CP requires copper atoms for functional maturation, and Ni²⁺, which shares coordination chemistry similar to Cu²⁺, can displace copper from nascent ceruloplasmin during hepatic synthesis, producing unstable apo-ceruloplasmin that is rapidly degraded. Differential detection analysis revealed markedly increased STEAP3 (Six-Transmembrane Epithelial Antigen of the Prostate 3) in the high-contamination group in the blood cell pellet. STEAP3 is a metalloreductase that converts transferrin-delivered Fe³⁺ back to Fe²⁺ within the endosome for cytosolic export via divalent metal transporter 1 (DMT1) and is the predominant STEAP family member in erythroid precursors and macrophages [55, 56]. The concurrent decrease in CP and increase in STEAP3 represents a coherent but maladaptive iron-handling signature. Extracellular Fe²⁺ accumulates because CP cannot oxidise it for transferrin loading, while cells compensate by upregulating STEAP3 to maximise iron extraction from the diminishing pool of properly loaded transferrin. This creates a metabolic paradox in which cells actively increase their iron-uptake machinery despite being in an iron-rich environment because the iron is in the wrong oxidation state. STEAP3 also possesses cupric reductase activity, and its upregulation has been associated with enhanced cellular copper uptake [57]. The additional roles of STEAP3 in p53-dependent apoptosis induction through interaction with proapoptotic protein Nix, and in cell cycle delay through positive regulation of Myt1 at the G2/M transition [56], suggest that its upregulation may also reflect a broader cellular stress response beyond iron compensation [58].

We identified a global decrease in actin-associated proteins in exposed birds. The Arp2/3 complex subunits ARPC4 and ACTR2 decreased in the high-contamination group; Arp2/3 is essential for branched actin network nucleation and plays a critical role in T-cell receptor signalling and cytoskeletal reorganisation during immune activation [59–62]. The actin regulator proteins formin-like-1 (FMNL1) and profilin (PFN1) were found to decrease in a dose-dependent manner in response to Cr. FMNL1 is selectively expressed in haematopoietic lineage cells [63], leukocytes [64], spleen and thymus [65], where it plays a role in cell adhesion, cytokinesis, cell polarization, and migration in mitosis [65]. FMNL1 regulates centrosome polarization in T cells, which is essential for the directed release of granules during cytotoxic T cell (CTL)-mediated killing, and depletion of FMNL1 in cytotoxic lymphocytes abrogated cell-mediated killing [62]. PFN1 inactivation produces multiple mitotic defects, including anaphase bridges, multipolar spindles, misaligned and lagging chromosomes, and failure of cytokinesis [66]. Actin is critical for lymphocyte activation, adhesion, and migration and the coordinated loss of these cytoskeletal regulators indicates that Ni and Cr toxicity fundamentally impair the capacity of blood cells to divide, migrate, and execute immune functions.

The most extensive perturbation, however, occurred within the nucleus. A dose-dependent decrease in nucleoporin NUP153 emerged as a central finding linking multiple downstream disruptions. NUP153 is an essential component of the nuclear basket that mediates nucleocytoplasmic transport of mRNA and ribonucleoprotein particles, maintains chromatin organisation at the nuclear periphery, docks splicing machinery to the nuclear pore, and acts as a global regulator of gene expression [67, 68]. Nuclear pore–chromatin interactions are increasingly recognised as regulators of gene expression, and loss of NUP153 has been shown to alter chromatin organisation and silence normal gene expression [69]. Critically, NUP153 contains four zinc finger motifs that bind Ran-GTP and Ran-GDP, and these motifs are essential for maintaining the GTP/GDP gradient across the nuclear envelope [70]. Because Ni²⁺ can substitute for Zn²⁺ in zinc-binding domains, we propose that nickel-mediated displacement of zinc from the zinc finger motifs of NUP153 destabilises its structure and function, providing a direct mechanistic link between nickel exposure and nuclear pore dysfunction.

The decrease in NUP153 was accompanied by reduced abundance of regulator of chromosome condensation 1 (RCC1), the sole guanine nucleotide exchange factor for Ran, which ensures that Ran-GTP is concentrated near chromatin while Ran-GDP predominates in the cytoplasm. The concurrent loss of NUP153 (which regulates Ran-GTP hydrolysis at the nuclear pore) and RCC1 (which generates Ran-GTP on chromatin) effectively dismantles both the “on” and “off” switches of the Ran cycle, producing catastrophic Ran-GTP depletion and loss of nuclear transport directionality. This dual failure destabilises not only nuclear–cytoplasmic transport but also DNA repair protein import, RNA export, chromatin condensation, and mitotic spindle formation, as well as other processes dependent on a functional Ran gradient. Consistent with this model, we observed a global decrease in histone proteins (H2A variants, H2B, H3.3A, H1.4), which may be both a cause and consequence of RCC1 decline, as RCC1 requires chromatin-bound histones for its localisation and activity [71].

The collapse of nuclear transport had predictable downstream consequences for RNA metabolism. Spliceosome and RNA processing components were broadly reduced, including the DEAD-box helicases DDX17 and DHX15, the splicing regulators RBMX, PRPF39, TRA2B, and RBM4B, the mRNA export factor ALYREF, and the exon junction complex component RNPS1. NUP153 directly docks splicing machinery to the nuclear pore, and its loss would impair both co-transcriptional splicing and mRNA export [68]. Histone dysregulation and post-translational modification changes resulting from heavy metal exposure likely compound this effect by reducing transcriptional activity at the chromatin level [72, 73], creating a convergent suppression of gene expression through both transcriptional repression and impaired mRNA maturation and export.

Against this backdrop of nuclear infrastructure collapse, vaccinia-related kinase 1 (VRK1) and thyroid hormone receptor interactor 13 (TRIP13) were both more abundant, representing a compensatory stress response to accumulating DNA damage and chromosomal instability likely arising from Cr-induced DNA damage [74, 75]. The upregulation of MTHFD1, a trifunctional enzyme in folate-mediated one-carbon metabolism that supplies nucleotide precursors for DNA synthesis and repair, further supports active engagement of the DNA damage response [76]. VRK1 is a nuclear serine/threonine kinase that phosphorylates histones H3 and H2A to drive chromosome condensation, controls chromatin relaxation associated with histone acetylation, and regulates the non-homologous end joining DNA repair pathway [72, 77, 78]. VRK1 activity is partially regulated through Ran-dependent nuclear localisation and interacts functionally with RCC1 [79], further linking its dysregulation to the collapse of Ran signalling. Crucially, VRK1 binds Ran-GDP for its own nuclear import; with the Ran gradient depleted, VRK1 protein may accumulate without achieving proper nuclear localisation or substrate access, representing a state of elevated abundance without corresponding activity. TRIP13, involved in cell cycle checkpoint control and DNA damage response, may facilitate checkpoint override that forces damaged cells through mitosis before repair is complete. The combination of active damage sensing (VRK1 increase) with potential checkpoint bypass (TRIP13 increase) creates a “repair trap” in which DNA damage response is initiated but cannot be completed, driving genomic instability [77, 80, 81]. Consistent with this, the key centromere and kinetochore proteins CENPC and CENPT were reduced, supporting defective chromosome segregation, which is a hallmark of weakened Ran gradient signalling and impaired RCC1 activity that normally coordinate spindle assembly and kinetochore function.

The upregulation of MYCBP2, an E3 ubiquitin ligase that ubiquitinates TSC2 to stimulate mTOR signalling and regulates apoptotic pathways, alongside the concurrent decrease in SQSTM1/p62, points toward heightened protein quality control under metal-induced proteotoxic stress. p62 is normally degraded during active autophagy, and its decline may indicate enhanced autophagic flux attempting to clear damaged histones and protein aggregates generated by heavy metal exposure. However, while the MYCBP2-mTOR axis and autophagy together represent maximal engagement of cellular clearance and survival pathways, they cannot restore the chromatin architecture and nuclear transport functions that have been dismantled upstream, leaving cells trapped between active damage clearance and irreversible structural collapse.

Together, these findings support a unified mechanism in which Ni and Cr toxicity targets the nuclear pore–Ran–chromatin axis, with NUP153 dysfunction acting as an upstream event that propagates global gene suppression, defective RNA metabolism, chromosomal instability, and failure of cellular stress-adaptation pathways in blood cells. Cytoskeletal integrity is tightly coupled to nuclear transport and chromatin architecture through the LINC complex and mechanical signalling pathways, and its collapse further amplifies transcriptional repression and cell cycle arrest. The epigenetic dimension of this toxicity, mediated by the proposed interaction of NUP153 with the histone acetyltransferases PCAF/p300 [82], may explain the well-documented gene-silencing effects of chronic nickel exposure, which have previously been attributed primarily to direct histone modifications but may in fact originate upstream at the nuclear pore alongside these modifications.

These molecular disruptions carry direct implications for avian fitness and population viability in contaminated environments. The coordinated suppression of immune-related proteins, including cytoskeletal regulators essential for leukocyte migration (PFN1, FMNL1, ACTR2), complement components (CFH, C9), and acute-phase proteins (CP, SAA1) indicates a state of functional immunosuppression that would reduce the capacity of exposed birds to mount effective responses against pathogens and parasites. The impairment of erythropoiesis, evidenced by decreased HEMGN and disrupted iron metabolism, suggests compromised oxygen-carrying capacity affecting aerobic performance, flight endurance, and thermoregulation, all which are traits critical for foraging efficiency and predator evasion in a free-living passerine. The disruption of cell division machinery through cytoskeletal collapse and centromere protein loss (CENPC, CENPT) may impair haematopoietic stem cell renewal, progressively depleting circulating immune and erythroid cell populations over time.

For a sedentary species inhabiting these ultramafic landscapes year-round, chronic exposure creates a cumulative physiological burden in which metal detoxification demands divert resources from reproduction, immune defence, and somatic maintenance. These findings suggest that house sparrows in heavily contaminated areas of New Caledonia are living in a state of subclinical multi-system compromise that, while not immediately lethal, may reduce survival, breeding success, and population persistence. New Caledonia’s ultramafic soils are naturally enriched in nickel and chromium, and decades of mining and smelting have raised these metals to even higher levels. Our results therefore raise a broader question: how have avian species adapted, or failed to adapt, to chronically metal-rich landscapes over evolutionary timescales?

## 5. Conclusion

This study provides the first integrated blood proteome characterisation of Ni and Cr toxicity in a free-living avian species, revealing a multi-system molecular cascade that connects nuclear pore dysfunction, chromatin collapse, DNA damage response failure, iron dyshomeostasis, cytoskeletal shutdown, and immunosuppression. The discovery that VAMS-based pellet proteomics provides substantially deeper proteome coverage than plasma alone establishes a new methodological framework for wildlife ecotoxicology and biomarker discovery. The identification of a three-metal interaction chain in which Ni disrupts Cu-dependent CP, triggering Fe dysregulation, and the proposed NUP153–Ran–chromatin axis as a unifying mechanism for Ni/Cr nuclear toxicity, represents novel mechanistic contributions that extend current understanding of heavy metal carcinogenesis and have translational relevance for human populations exposed to similar metal combinations in mining and industrial settings.

## 6. Acknowledgments

The authors acknowledge support from Janet Chik. Some of the research described herein was facilitated by access to the Australian Proteome Analysis Facility (APAF) funded under the Australian Government National Collaborative Research Infrastructure Strategy (NCRIS)/Education Investment Fund.

